# Partial Destabilization of Amyloid-β Protofibril by Methionine Photo-oxidation: A Molecular Dynamic Simulation Study

**DOI:** 10.1101/2022.05.29.493920

**Authors:** Fahimeh Maghsoodi, Tye D. Martin, Eva Y. Chi

## Abstract

Selective photosensitized oxidation of amyloid protein aggregates is being investigated as a possible therapeutic strategy for treating Alzheimer’s disease (AD). Photo-oxidation has been shown to lead to the degradation of amyloid-β (Aβ) aggregates, ameliorate aggregate toxicity and reduce aggregate levels in the brains of AD animal models. To shed light on the mechanism by which photo-oxidation induces fibril destabilization, we carried out an all-atom molecular dynamics (MD) simulation to examine the effect of methionine (Met35) oxidation on the conformation and stability of a highly β-sheet-rich Aβ_9-40_ protofibril composed of 12 monomers. Analyses of up to 1 μs MD simulations showed that the oxidation of the Met35 residues reduced the overall conformational stability of the protofibril. Specifically, Met35 oxidation that resulted in the addition of hydrophilic oxygens disrupted the hydrophobic interface that stabilizes the stacking of the two hexamers that composed the protofibril. The oxidized protofibril is more solvent exposed, less compact, and exhibits more backbone flexibility. However, it retained the underlying U-shaped architecture of each peptide. Although some loss of β-sheets occurred, a significant portion (~76%) remained. twisting of the peptides along the protofibril axis was observed, and the hexamers remained. Our simulation results are thus consistent with our experimental observation that photo-oxidation of Aβ40 fibrils results in the dis-agglomeration and fragmentation of Aβ fibrils, but did not cause complete disruption of the fibrillar morphology or β-sheet structures. The partial destabilization of Aβ aggregates supports the further development of photosensitized platforms for the selective targeting and clearance of Aβ aggregates as a therapeutic strategy for treating AD.

## Introduction

The abnormal aggregation and deposition of the amyloid-beta (Aβ) peptides into extracellular amyloid plaques is a major pathological event in the development of Alzheimer’s disease (AD).^1–3^ It is linked to the progressive neurodegeneration in AD^4–8^ that involves the impairment of synaptic transmission and the loss of long-term potentiation.^9,10^ Amyloid plaques are formed by the misfolding and aggregation of Aβ into small oligomers that subsequently grow into large fibrils rich in cross β-sheets.^11,12^ Aβ oligomers have been found to be cytotoxic as their interactions with neuronal cells cause increases in cell membrane conductance and calcium influx which in turn lead to neuronal apoptosis.^13–16^ Aβ fibrils also contribute to neurodegeneration by impairing axonal transport^10,13^ and seeding the aggregation of the tau protein to form intracellular neurofibrillary tangles.^14^ Additionally, Aβ aggregates are also involved in the spatiotemporal disease progression through cell-to-cell transmission.^15–17^ Because of the central roles Aβ aggregates play in AD pathogenesis, modulating the peptide’s aggregation and inducing the selective degradation and clearance of Aβ aggregates is an attractive therapeutic strategy.

Among the many approaches that have been studied, photodynamic therapy (PDT) has drawn the attention of researchers^18–20^ because it is spatiotemporally controllable and minimally invasive.^21,22^ PDT has been used in to treat diseases since the 1960s^23^ and is currently being used to treat many types of skin, lung, and esophageal cancers or pre-cancers.^24^ PDT requires a photosensitizer which is excited into a triplet state upon light exposure and causes the formation of singlet oxygens, which then leads to the generation of reactive oxygen species (ROS). ROS can then oxidize cellular components, including cell membranes and organelles, and induce apoptosis, which destroys diseased tissues.^25–27^ In recent years, a number of studies have investigated whether PDT can be a promising therapeutic strategy for treating AD. A range of photosensitizers have been tested and these studies have shown that photo-oxidation of monomeric (or soluble) Aβ can inhibit the peptide’s aggregation and that photo-oxidation of fibrillar Aβ can cause fibril fragmentation and disintegration.^18,28-36^ Encouragingly, Aβ aggregate degradation induced by photo-oxidation has also be found to attenuated aggregate toxicity^37^ and reduced aggregate levels in the brains of AD mouse models.^38^ In transgenic AD models of *C. elegans*, photo-oxidation of Aβ fibrils by fullerene based up-conversion nanoparticles^39^ and porphyrin-based metal-organic frameworks^28^ have been found to reduce Aβ neurotoxicity and extend the longevity of *C. elegans*.

Most of the compounds that have been studied are known photosensitizers and are non-selective, including polyoxometalate^29^, 1,2,4-oxadiazole^33^, tetra(4-sulfonatophenyl) porphyrin^32^, rose bengal^34^, methylene blue^36^, and porphyrinic metal-organic frameworks^28^, and induce the photooxidation of both Aβ monomers and aggregates, as well as other biomolecules in the vicinity of the photosensitizers. This causes off-target oxidation and is a major drawback of PDT.^40,41^ Recently, several aggregate-selective photosensitizers have been developed and tested, including those based on fibril-binding dyes thioflavin-T and curcumin^37,38,42^ and amyloid aggregate selective *p*-phenylene ethynylene-based florescence sensors.^43–45^ These new aggregate-selective photosensitizers can potentially overcome off-target oxidation and minimize side effects in future clinical applications.

To further develop photoactive platforms that target the degradation and clearance of Aβ amyloids, a fundamental understanding of the effect of photo-oxidation on Aβ aggregates is needed. Many studies have documented the morphological changes to the fibrillar structure of Aβ upon photo-oxidation, including fibril fragmentation, rupture, and disintegration.^36,37,43^ Photooxidation sites have also been identified. However, molecular level details of the conformational changes that leads to fibril destabilization have not been elucidated.

Several studies have shed light on the mechanism by which photo-oxidation attenuates the aggregation of Aβ monomers. For Aβ_1-42_ monomers, thioflavin T sensitized the oxidation of Tyr10, His13, His14, and Met35 residues^46^, which reduced the aggregation propensity of the monomers and delayed the formation of both oligomers and protofibrils. Oxidized Aβ_1-42_ monomers also did not exert any neurotoxicity^47^, which may be explained by the finding that Aβ with oxidized Met35 did not penetrate the plasma membrane of neurons and when they were extracellularly applied.^48^ An NMR study demonstrated that oxidation of Met35 in Aβ monomers considerably impeded aggregation and the propensity β-strand formation as the addition of a hydrophilic oxygen disrupted the hydrophobic interactions that stabilize the β-strands.^49^ These findings are consistent with others that found oxidation of Met35 inhibited coil-to-β-sheet transition^50^, significantly reduced trimer and tetramer formation,^51^ and slowed the rate of fibrillation.^52^ In an *in vivo* experiment, oxidation of Met35 in Aβ Aβ_1-42_ prevented the formation of a paranucleus^53^ and thereby inhibiting further fibrillation. Computational studies corroborated experimental findings and indicated that the oxidation of Met35 in Aβ impeded aggregation by reducing β-strand content on the C-terminal hydrophobic region.^54^

For Aβ_1-40_ fibrils, we and others have shown that photosensitization results in the oxidation of two histidine residues (His13 and His14) and Met35.^18,43,46^ His13 and His14 are located on the surface of Aβ fibrils whereas Met35 is located in the hydrophobic core region of the fibrils. Concomitant with photo-oxidation, we observed that long fibrils that formed clumps disaggregated and fragmented into shorter fibrils by AFM imaging. The shorter oxidized fibrils largely retained the β-sheet structures that were present in the native fibrils and the ability to seed fibrillation of Aβ monomers.^43^ Photo-oxidation of Aβ fibrils thus destabilized some structural features of Aβ fibrils, but did not completely disassemble the aggregates. Here, we carried out an all-atom molecular dynamics (MD) study to gain an understanding of the effect of oxidation on fibril conformational dynamics and stability and focused our analysis on the oxidation of Met35 as it is located in the fibril core and has been found to significantly attenuate the assembly of Aβ monomers into ordered aggregates. MD simulation has been widely used as a powerful tool to study the self-assembly of biomolecules and gain deeper insights into biomolecular interactions and structural dynamics.^55–59^ For this study, MD can reveal early structural changes to the extended β-sheet conformation of Aβ fibrils that results from Met35 oxidation, which ultimately lead to fibril destabilization. All atom MD simulations for up to 1 μs were carried out on a β-sheet rich Aβ_9-40_ protofibril. A number of analytical tools were used to assess the effect of Met35 oxidation on protofibril conformation and dynamics. Specifically, we monitored the deviation of the protofibril from its initial structure and analyzed changes to secondary structures that stabilize the protofibril.^60,61,62,63^ Global changes to the protofibril conformation were also monitored through analyzing the number of hydrogen bonds and solvent accessible surface area. Overall, we sought to gain an understanding of the dynamic destabilization of an Aβ protofibril after substituting Met35 with its oxidized form and reveal the underlying mechanism of photo-oxidation induced partial fibril destabilization.

## Methods

### System setup

The initial structure of the Aβ protofilament (Aβ_9-40_) was obtained from the Protein Data Bank (PDB ID: 2LMN), which consisted of two stacked hexamers.^64^ Peptide chains are designated from A to F for the top hexamer and G to L for the bottom hexamer (Figure 1A). The methionine 35 (Met35) residues are colored yellow; they are located in the middle of the protofibril where the hydrophobic surfaces of the two hexamers meet (Figure 1A). The structures of Met and Met^ox^ are shown in Figure 1B. Protofibrils containing either Met35 (Aβ_9-40_-Met35) or oxidized Met35 (Aβ_9-40_-Met35^ox^) were set up. All-atom, explicit solvent molecular dynamics (MD) simulations were carried out for the native and oxidized protofibril systems for 300 ns in triplicates. Additionally, one of the 300 ns simulation for each protofibril system was carried out to a longer simulation time of 1 μs.

**Figure 1.**
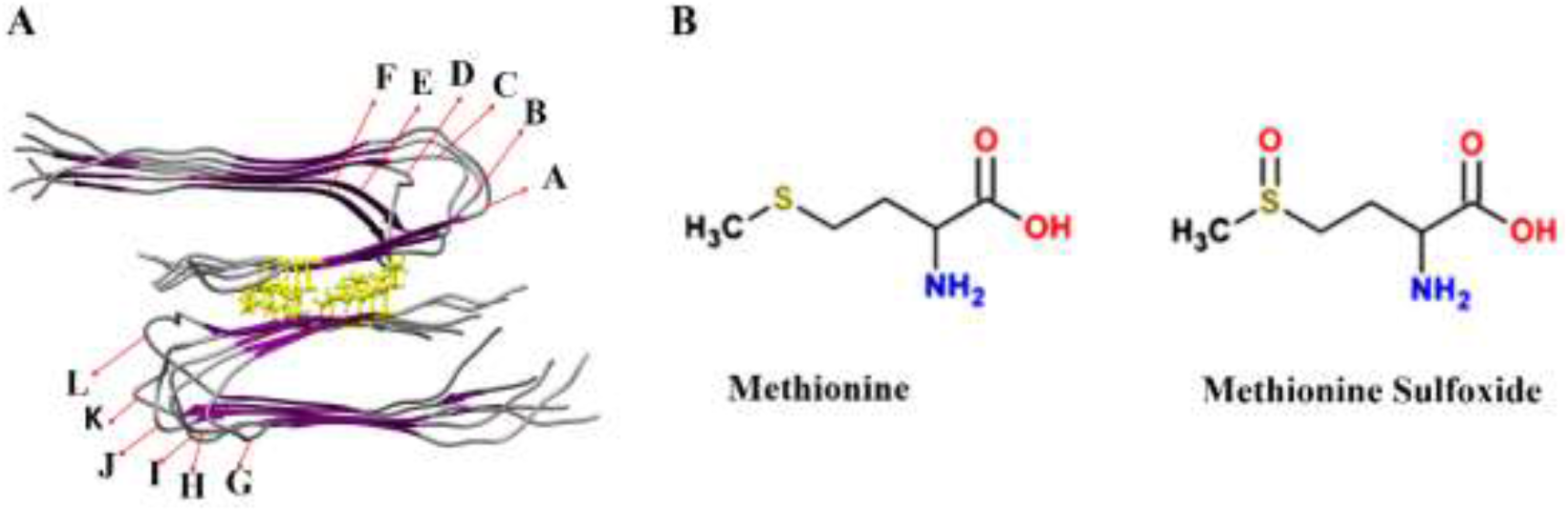
**A**: Structure of Aβ protofibril (2LMN) which consists of two hexamers. Individual Aβ_9-40_ chains are denoted as A to L and Met35 residue side chains are shown in yellow. Purple and gray in the protofibril represent β-sheet and coil structures, respectively. **B**: Chemical structures of Met and Met^ox^.

### Simulation methods

MD simulations were prepared and run using Gromacs v2018^65^ in an isothermal (NVT) and isobaric ensemble (NPT) using the CHARMM27 force field and the TIP3P water model. The initial structure and parameterization of Met35^ox^ were obtained using SWISS-PARAM, which has been used in studies investigating the interactions of proteins including Aβ with small organic molecules^66^. To prepare the oxidized protofibril system (Aβ_9-40_-Met35^ox^), each of the 12 Met35 residues in the native Aβ protofibril was substituted with a Met35^ox^. A cubic simulation box of 10.8 nm in each dimension with periodic boundary conditions was used for the native and oxidized Aβ protofibrils. Sodium ions were then added to maintain charge balance.

Energy minimization of the systems was carried out using the steepest descent minimization method to reach a maximum force < 1000.0 kJ/mol/nm. The energy minimized systems were then equilibrated for 1 ns in NVT and NPT ensembles using LINCS^67^ constraints for H-bonds with a standard leap-frog integrator selected with a 2 fs time step. Van der Waals interactions were treated with a cutoff of 1.0 nm and long-range electrostatic interactions were calculated using Particle Mesh Ewald (PME) with a 1.0 nm cutoff length.^68^ For NVT equilibration, the modified Berendsen thermostat coupling was used to keep the temperature constant at 310 K. For NPT equilibration, the temperature (310 K) and pressure (1 bar) coupling was applied *via* the modified Berendsen thermostat coupling^69^ and Parrinello–Rahman method^70^, respectively. MD productions of equilibrated systems were performed at NPT. For each system, three trajectories of 300 ns were run, and averaged results are reported. One randomly selected trajectory of each system continued to run to 1 μs. UCSF chimera^71^ was used for visualization and further analyses were done using Gromacs v2018. All simulations were run on the Comet hybrid computing cluster at the Extreme Science and Engineering Discovery Environment (XSEDE) digital service at the San Diego Supercomputer Center (SDSC).^72^

### Analysis of simulation results

A number of different tools provided by the Gromacs v2018 package^65^ were utilized to analyze the simulation results which focused on assessing the structural stability of the protofibrils. The evaluated parameters include root mean square deviation (RMSD), root mean square fluctuation (RMSF), radius of gyration (R_*g*_), solvent accessible surface area (SASA), total number of hydrogen bonds (H-bonds), and intra- and inter-chain Asp23 (D23) - Lys28 (K28) salt bridges. Moreover, the secondary structures of the protofibrils were investigated *via* the Dictionary of Secondary Structure of Protein (DSSP)^73^ program to assess any conformational changes to the protofibril with Met35^ox^ substitutions.

RMSD was obtained based on the C_α_ atoms of the peptide chains for the two hexamers. The R_*g*_ values of native and oxidized Aβ protofibrils were computed using the *gmx gyrate* tool. The GROMACS utility *gmx bond* was used to compute the number of H-bonds between all main chains of the protofibril where the donor-acceptor cut-off distance was assigned to be 0.35 nm. SASA values of the protofibrils were calculated for all simulations. In addition, SASA values of the Met35^ox^ and Met35 residues were calculated for the 1 μs simulation trajectories. The distance between Met35 residues of oppsing chains (A and G, B and H, C and I, D and J, E and K, F and L) in native and oxidized protofibrils were determined and compared. *GMC mak_ndx* was used to create appropriate index files separating Met35 residues of opposing Aβ chain pairs for all simulation trajectories. Salt-bridge is another important contributor to protein stability.^74,75^ In the protofibrils, D23-K28 distances on the same peptide chains and between adjacent chains for all simulations were determined.

### Statistical analysis

Three trajectories of the 300 ns simulations were carried out. Averaged values of RMSD, R_*g*_, SASA, RMSF, Met35-Met35 and Met35^ox^-Met35^ox^ distances, D23-K28 salt bridge distances, and the total number of H-bonds were calculated and reported. *P*-values were calculated using a paired *t*-test. To compare the mean values for a specific analysis, for example, R_*g*_ values of native vs. oxidized protofibril, one-tailed paired *t*-test with a null hypothesis (means are identical) and an alternative hypothesis (value obtained from oxidized Aβ protofibril is larger than that from the native protofibril) was carried out at a significance level of 0.05.

## Results

The fibrillar conformation of the Aβ peptide is highly thermodynamically stable and is primarily stabilized by the extended β-sheet core. We and others have shown that photosensitizer-mediated oxidation of Aβ fibrils results in the oxidation of His13 and His14, which are located on the surface of the fibrils, and Met35, which is located in the β-sheet core of the fibrils^18,43,46^. We observed that photo-oxidation caused clumps of long fibrils to disaggregate and fragment into short fibrils. These shorter oxidized fibrils largely retained the β-sheet structures that were present in the native fibrils and the ability to seed fibrillation of Aβ monomers.^43^ To gain a molecular-level understanding of the effect of oxidation on fibril structure and stability, we focused our analysis on the oxidation of Met35 as it is located in the fibril core and is expected to have the most significant impact on fibril structure and stability. An Aβ_9-40_ protofibril with two stacked hexamers was chosen in this study as it contains the salient structural features of Aβ amyloid polymorphs, the U-shaped architecture of each Aβ peptide, extended inter-sheet chain-chain contacts, and hydrophobic strand-strand contacts.^76^ We first examined snapshots of native and oxidized protofibril trajectories. Then, we calculated RMSD, R_*g*_, SASA, and RMSF values to evaluate structural fluctuations and possible expansion of the protofibril. Secondary structures, the number of H-bonds, and salt bridge interactions were also analyzed to gain more insights into the effects of Met35 oxidation.

### Dynamics of native and oxidized Aβ protofibrils

Snapshots of native (Aβ_40_-Met35) and oxidized (Aβ_40_-Met35^ox^) Aβ protofibrils from the 1 μs simulations are shown in Figure 2. Snapshots from the 300 ns trajectories are shown in Figure S1 of Supplemental Information. As shown in Figure 2, both protofibrils undergo structural changes with simulation time and become less ordered compared to the energy minimized structures at beginning of the simulations (0 ns). The extended β-sheets started to twist along the protofibril axis such that some of the β-strands are no longer co-planar by the end of the simulations. Chains located at the edges of the protofibrils appeared to have moved the most.

**Figure 2.**
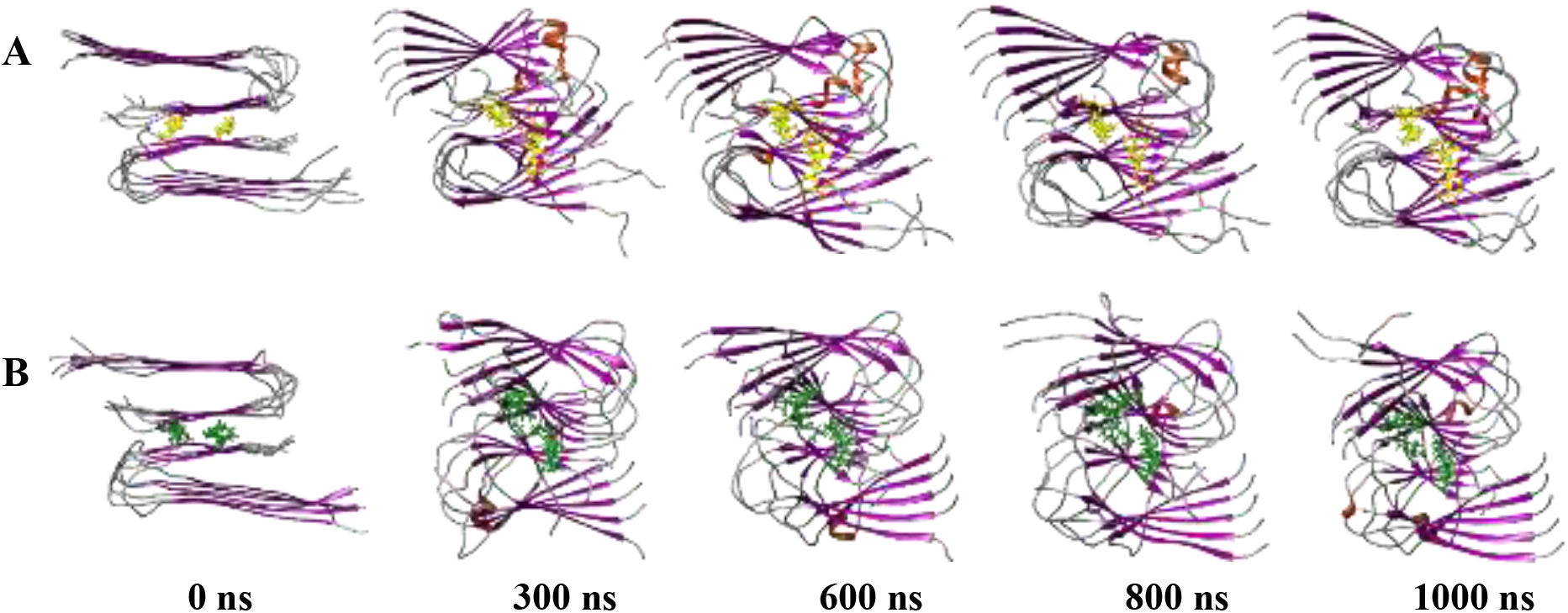
Snapshots of native (**A**) and oxidized (**B**) Aβ protofibrils from 1 μs simulations. Met35 and Met35^ox^ are shown in yellow and green, respectively. β-strands, α-helices, and coils are shown in purple, orange, and grey, respectively.

To better visualize chain movements and compare the extents of chain twisting between Aβ_40_-Met35 and Aβ_40_-Met35^ox^ protofibrils, the end chains (A and L) in both protofibrils were specifically highlighted and compared side by side at the end of the 1 μs simulations (Figure 3). As shown, chain A (magenta) in the native protofibril is twisted but largely retained its β-hairpin structure (Figure 3A) whereas the same chain in the oxidized protofibril is also twisted but the β-hairpin is less aligned (Figure 3C). At the other end of the protofibril, chain L (blue) in the native protofibril is twisted along the protofibril axis, lost some of its β-sheet characteristics, and one end of the chain adopted a short helical structure (Figure 3B). In comparison, chain L in the oxidized protofibril is significantly more twisted along protofibril axis, by about 90° (Figure 3D). Although this chain largely retained its hairpin structure, intra-chain β-sheet contacts are lost as well as the inter-chain β-sheet contacts with the adjacent chain K in the protofibril. The turn region also adopted an α-helical structure.

**Figure 3.**
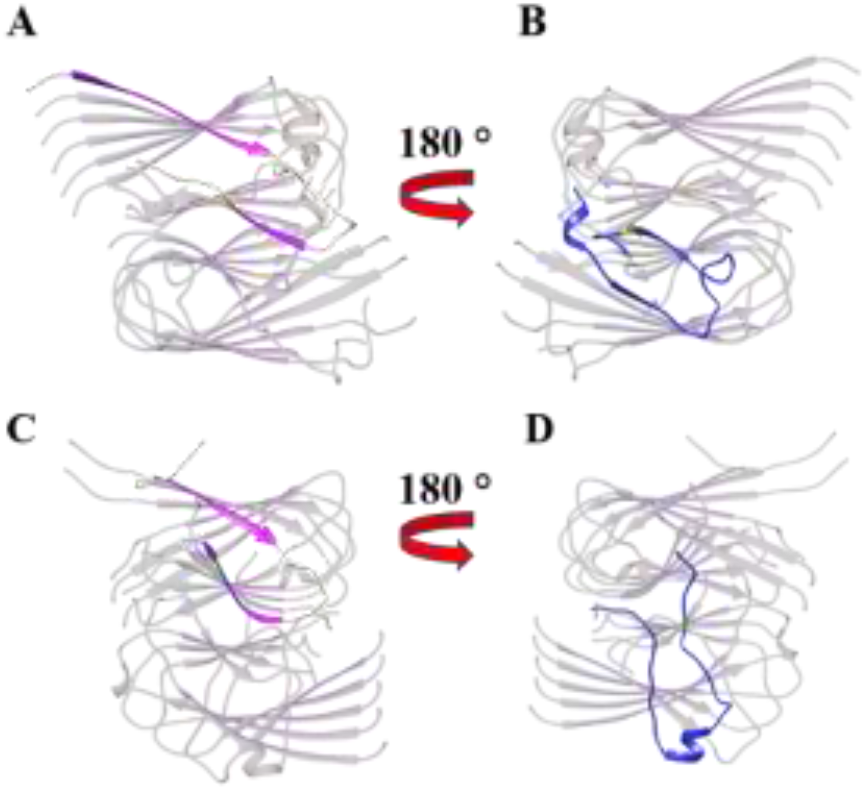
Visualization of end chains A (magenta) and L (blue) of the native (**A** and **B**) and the oxidized (**C** and **D**) Aβ protofibrils at 1000 ns of simulation time. Structures in **A** and **C** are each rotated 180° to visualize the other ends of the protofibrils. The rest of the chains are colored gray for better visualization of the end chains.

Overall, by visual inspection, both protofibrils largely retained their aggregated structures. Although the difference between Aβ_40_-Met35 and Aβ_40_-Met35^ox^ protofibrils in these snapshots are not large, the twisting of chains was observed more frequently and at a higher degree in the Aβ_40_-Met35^ox^ protofibril compared to the Aβ_40_-Met35 protofibril. Structural loss is also more apparent in the oxidized protofibril.

### Structural stability of native and oxidized Aβ protofibrils

To assess the global conformational stabilities of the two protofibrils, we performed a number of analyses, including calculating the Cα-RMSD, R_*g*_, SASA, and RMSF values of the 300 ns and 1 μs trajectories. RMSD values, measured with respect to the initial energy minimized structures, of the short (300 ns) and long (1 μs) simulations are shown in Figure 4A and 4B, respectively; values in Figure 4A are averages from three 300 ns trajectories. The RMSD values of the native protofibril stabilized to about 0.62 nm after a sharp initial increase. RMSD values for the oxidized protofibril were higher than those of the native protofibril and reached above 0.8 nm at the end of the 1 μs simulation (Figure 4B). Moreover, the RMSD profile for the oxidized protofibril showed more fluctuations and rather than leveling off, showed an increasing trend. The higher RMSD values, along with greater fluctuations and an increasing trend, indicate a destabilizing effect of Met35 oxidation on the protofibril.

**Figure 4.**
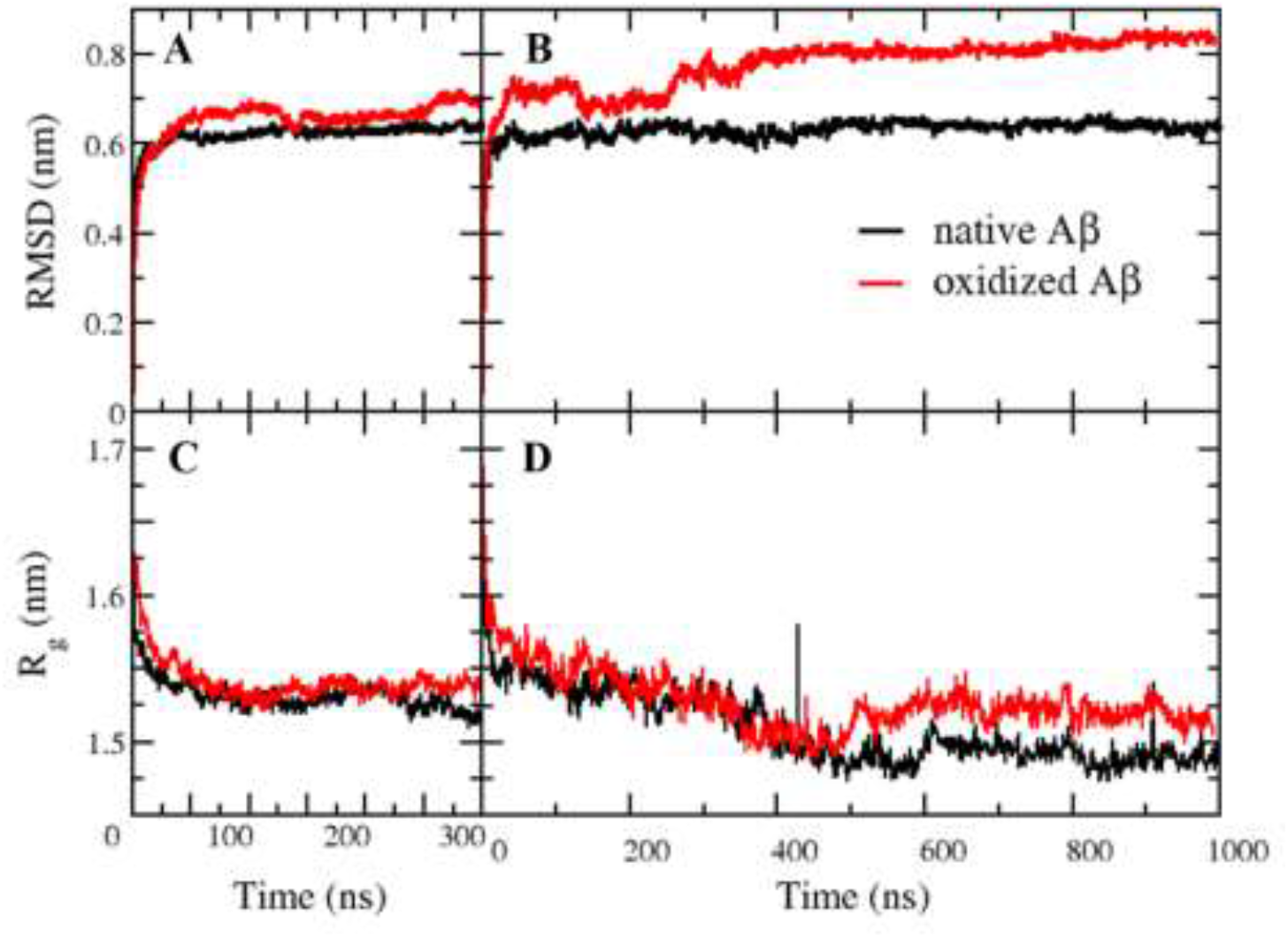
RMSD (**A** and **B**) and R_*g*_ (**C** and **D**) plots of native (black) and oxidized (red) protofibrils from 300 ns (A and C) and 1 μs (B and D) simulations. RMSD and R_*g*_ values shown for the 300 ns simulations (A and C) are averages from 3 simulation trajectories.

The radius of gyration (R_*g*_) of the native and oxidized protofibrils over the 300 ns and 1 μs trajectories are shown in Figure 4C and 4D, respectively. R_*g*_ values are calculated from the mass-weighted spatial distribution of the atoms in the protofibril and can be interpreted as a measure of the structural compactness of the protofibril. Both native and oxidized protofibrils exhibited similar R_*g*_ values of around 1.52 during the 300 ns trajectories (Figure 4C). After 400 ns of simulation however, R_*g*_ values for the oxidized protofibril were slightly higher than those of the native protofibril (Figure 4D). Although the difference is small, it is statistically significant (p < 0.05) implying that Met35 oxidation moderately decreased the compactness of the protofibril, perhaps through disrupting the packing of the highly ordered and dense fibril core.

Solvent accessible surface area (SASA) is another important property that gives information about the overall protein conformation in an aqueous environment. Proteins are composed of hydrophobic and hydrophilic residues and tend to adopt structures that minimize the exposure of hydrophobic residues to the aqueous solvent. Increases in SASA from a stable state can indicate protein instability, such as unfolding that exposes hydrophobic residues to the solvent, which can lead to further undesirable changes such as irreversible aggregation. Substitution of amino acids, whether mutational or chemical, can also disturb the native conformation of a protein and result in partial unfolding which leads to increases in SASA.

In this study, SASA values for the protofibrils were computed for the 300 ns and 1 μs simulation trajectories. As shown in Figure 5A and 5B, SASA values of the oxidized protofibril were larger than those of the native protofibril; the average SASA values computed from the three 300 ns simulations for the native and oxidized Aβ protofibrils were 188 and 197 nm^2^, respectively. Met35 oxidation thus caused about 9 nm^2^, or 5% increase of protofibril SASA. For the 1 μs simulation, average SASA values of native and oxidized Aβ protofibrils were found to be 175 and 200 nm^2^, respectively. Although the SASA increase of 25 nm^2^ from the 1 μs simulation is larger than the 9 nm^2^ obtained from the 300 nm simulations, both show higher SASA values for the oxidized protofibril. To assess the effect of oxidation on solvent exposure of the Met35 residue, SASA values of the Met35 or Met35^ox^ residues for all simulation trajectories were also calculated and plotted (Figure 5C and 5D). As shown, Met35^ox^, located in the core of the protofibril, showed higher (by about 2 nm^2^ or 10%) SASA values compared to the Met35 in the native protofibril. Oxidation of Met to methionine sulfoxide increases the hydrophilicity of the residue and even though Met is deeply buried in the hydrophobic core of the protofibril, its solvent exposure increased. However, the ~2 nm^2^ increase in Met35 SASA only partially contribute to the overall 9-25 nm^2^ increase in protofibril SASA, indicating that oxidation of the Met35 at the core of the protofibril caused global conformational changes to the protofibril such that the oxidized structure is in a more solvent exposed state.

**Figure 5.**
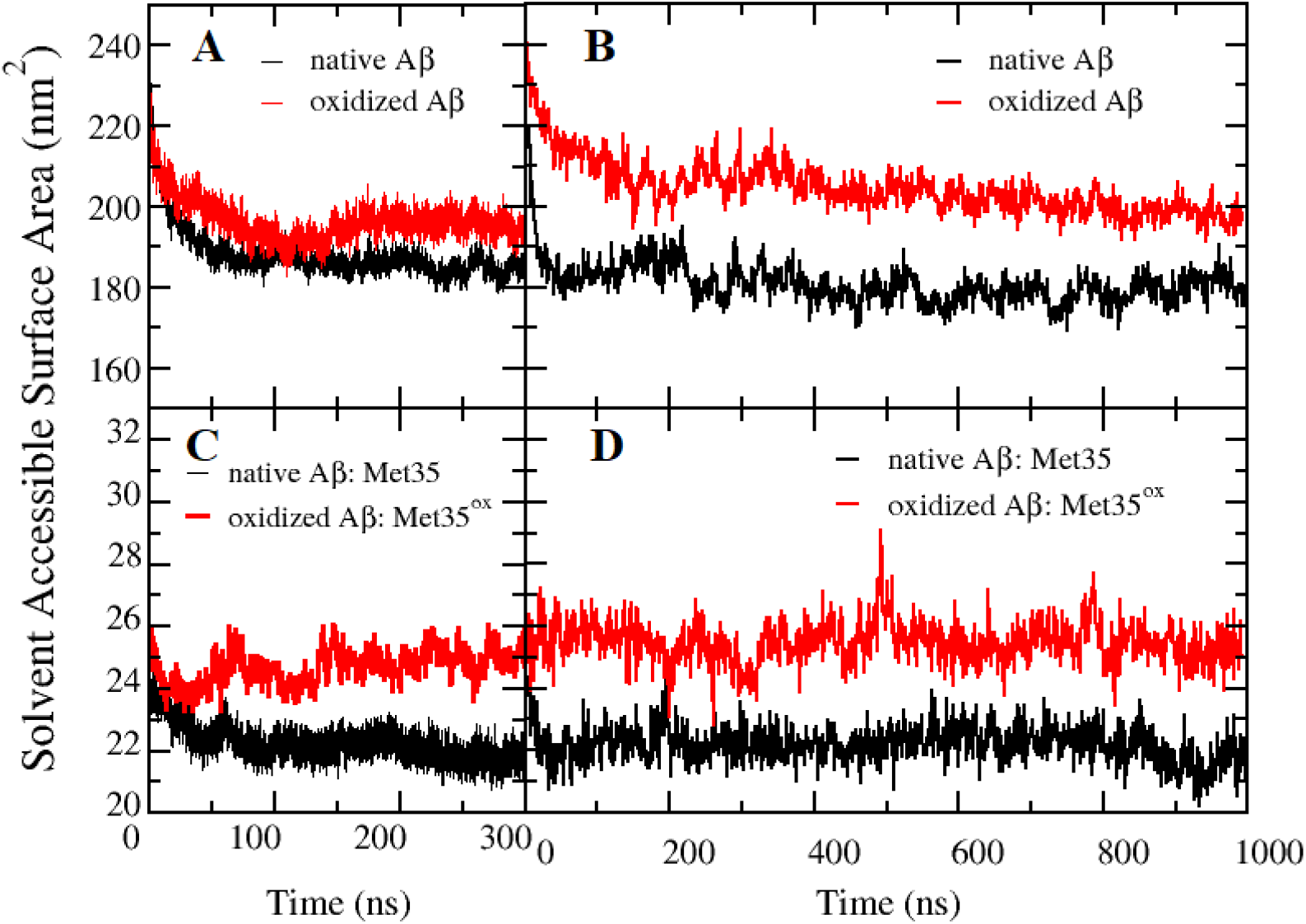
Solvent accessible surface area (SASA) values of the native (black) and oxidized (red) protofibrils calculated from 300 ns (**A**) and 1 μs (**B**) simulation trajectories. SASA values of the Met35 (black) or Met35^ox^ (red) residue in native and oxidized Aβ protofibrils calculated from 300 ns (**C**) and 1 μs (**D**) simulations.

To assess the local dynamics and flexibility of each residue of the Aβ chains, the RMSF values of the backbone of Aβ in the top hexamer (chains A to F) and bottom hexamer (chains G to L) for native and oxidized protofibrils after 1 μs of simulation were calculated and plotted in Figure 6. RMSF plots of the 300 ns simulations are shown in Figure S2 of the Supplemental Information. As expected, terminal chains A and F of the top hexamer and chains G and L of the bottom hexamer showed higher fluctuations with higher RMSF values than the interior chains (Figure 6A and 6C). Moreover, in the oxidized protofibril, 8 out of 12 chains (chains A, E, F, G, I, J, K, and L) showed statistically significant higher RMSF values compared to the native protofibril (p < 0.05). Some of these chains are terminal chains and some are in the interior of the protofibril. As such, our results show that Met35 oxidation caused increased chain flexibilities throughout the protofibril.

**Figure 6.**
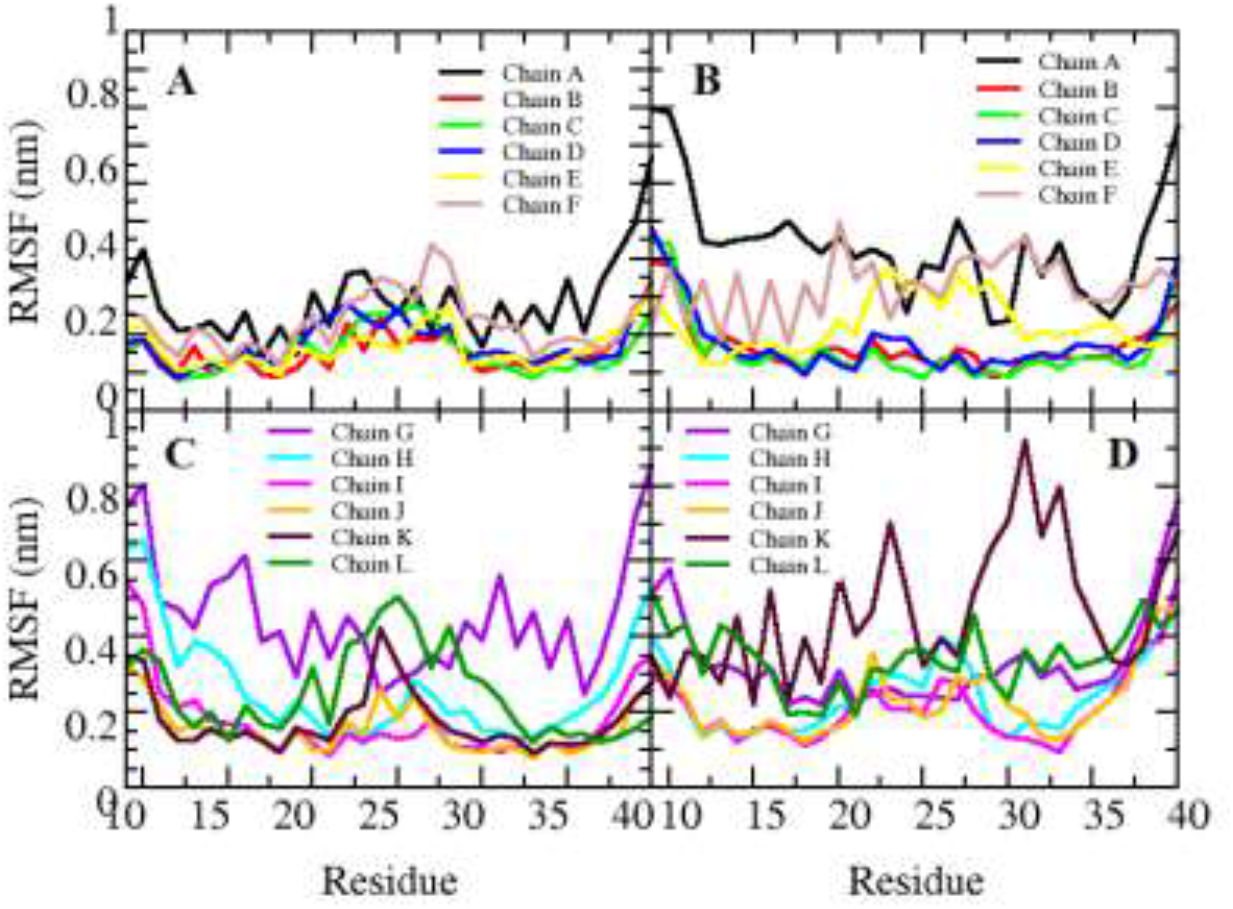
RMSF plots of different Aβ chains belonging to the top hexamer (chains A to F: **A** and **B**) and bottom hexamer (chains G to L: **C** and **D**) of the native (**A** and **C**) and oxidized (**B** and **D**) protofibrils after 1 μs of simulation.

Taken together, our analyses of the global conformational characteristics of a native Aβ_40_-Met35 and an oxidized Aβ_40_-Met35^ox^ protofibrils indicate that Met35 oxidation has a destabilizing effect on the highly ordered protofibril structure, wherein the Aβ_40_-Met35^ox^ protofibril showed higher values of RMSD, R_*g*_, SASA, and RMSF compared to those of the Aβ_40_-Met35 protofibril. Both protofibrils deviated from the initial energy minimized structures wherein twisting and misalignment of the β-strands were observed after 1 μs of simulation. The oxidized protofibril showed an approximately 30% increase in RMSD, 5-14% increase in SASA, and a smaller around 3% increase in R_*g*_. Compared with the native Aβ_40_-Met35 protofibril, 8 out of 12 chains of the oxidized protofibril showed higher RMSF values, or structural fluctuations, compared to the native protofibril.

### Methionine-Methionine distances in the protofibril core

The hydrophobic interactions of Met35 residues from the two opposing hexamers facilitate the favorable interaction of the two hexamers in the native protofibril (Figure 1). This parallel transversal combination of the two hexamers around the longitudinal axis of the protofibril is a common feature of various segmental polymorphisms identified in the Aβ fibril architecture^77^ and serves to shield hydrophobic residues at the C-termini of the Aβ peptides (Figure 1). Theoretical and experimental work have shown that residues Ile31, Met35, and Val39 are involved in the hydrophobic interface of both Aβ40 and Aβ42 fibrils.^76,78-81^

Because of the important role Met35 plays in stabilizing the fibril structure, we analyzed the packing of the Met35 residues in the protofibril core by measuring the distances between Met35-Met35 residues in the native protofibril and between Met35^ox^-Met35^ox^ residues in the oxidized protofibrils; results from the 1 μs simulations are summarized in Figure 7. The same plots for the 300 ns simulations are shown in Figure S3 of Supplemental Information. As shown in Figure 7, except for the end chains A and G, all other Met35-Met35 pairs equilibrated to 0.8 to 1 nm distances in the native protofibril. The end chains A and G showed greater Met35-Met35 distances of around 1.8 nm during most of the simulation and dropped to around 1 nm after 800 ns of simulation. This greater fluctuation is consistent with higher RMSF values observed for the two chains (Figure 6A and 6C). Met35^ox^-Met35^ox^ distances in oxidized fibrils in general showed more fluctuations during the simulation. In particular, interior chain pairs, C and I, D and J, and E and K, showed statistically significant higher Met35^ox^-Met35^ox^ distances than Met35-Met35 distances in the native protofibril (p < 0.05), while the end chain pairs (A and G, F and L) and the near end-chain pair (B and H) did not show significantly differences in Met-Met distances.

**Figure 7.**
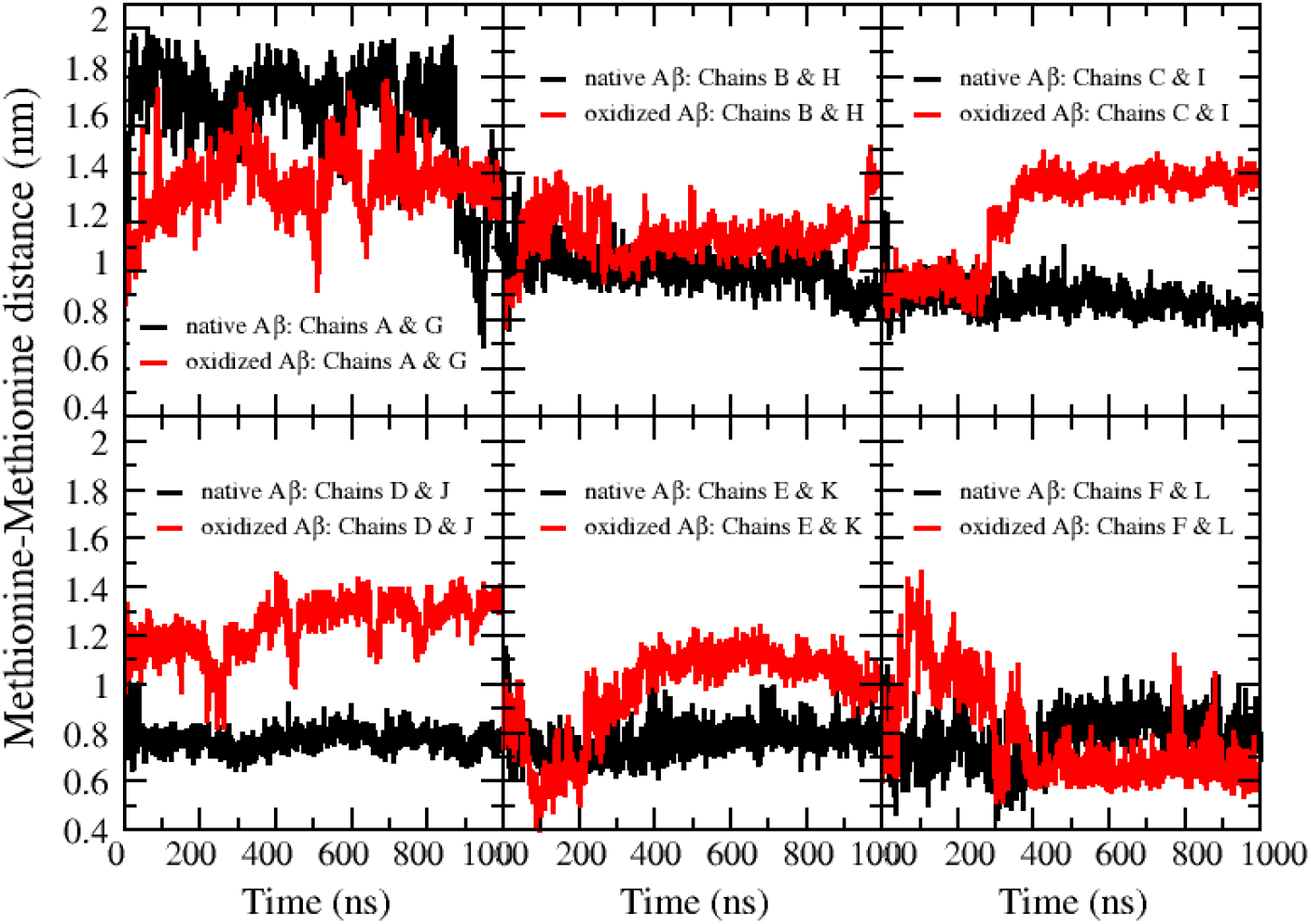
Distances between Met35-Met35 (black) and Met35^ox^-Met35^ox^ (red) residues of 6 opposing Aβ chain pairs (A and G, B and H, C and I, D and J, E and K, and F and L) in the protofibrils for the 1 μs simulation.

The trends in methionine-methionine distances suggest that the oxidation of the most hydrophobic or buried methionine residues of the central chains away from protofibril ends had the biggest impact in disrupting fibril structure. This finding also suggests that the increase in SASA of the protofibril with oxidation (Figure 5) is due to increased solvent exposure of the interior Met35 residues upon oxidation.

### Intra- and inter-peptide salt bridge distances in protofibrils

The Asp23-Lys29 (D23–K28) salt bridge that forms near the turn region of the Aβ peptide plays an important role in stabilizing the U-shape β-strand-turn-β-strand motif of the peptide and prevents large backbone motion. We performed an analysis of the D23–K28 salt bridge distances in the native and oxidized protofibrils to assess if Met35 oxidation adversely affected the stability of the intra-peptide salt bridges. D23–K28 distance for each Aβ chain for the 1 μs simulations are shown in Figure 8. As shown, comparisons between salt bridge distances of the native protofibril and oxidized fibrils did not show a consistent trend. Distances are comparable for chains E, G, H, I, and L. They are larger in the oxidized protofibril than the native protofibril for chains J and K, but smaller for chains A, C, and D. Thus overall, Met35 oxidation did not have a consistent effect on intra-molecular salt bridge distances, and these salt bridges appear to be relatively unaffected by Met35 oxidation.

**Figure 8.**
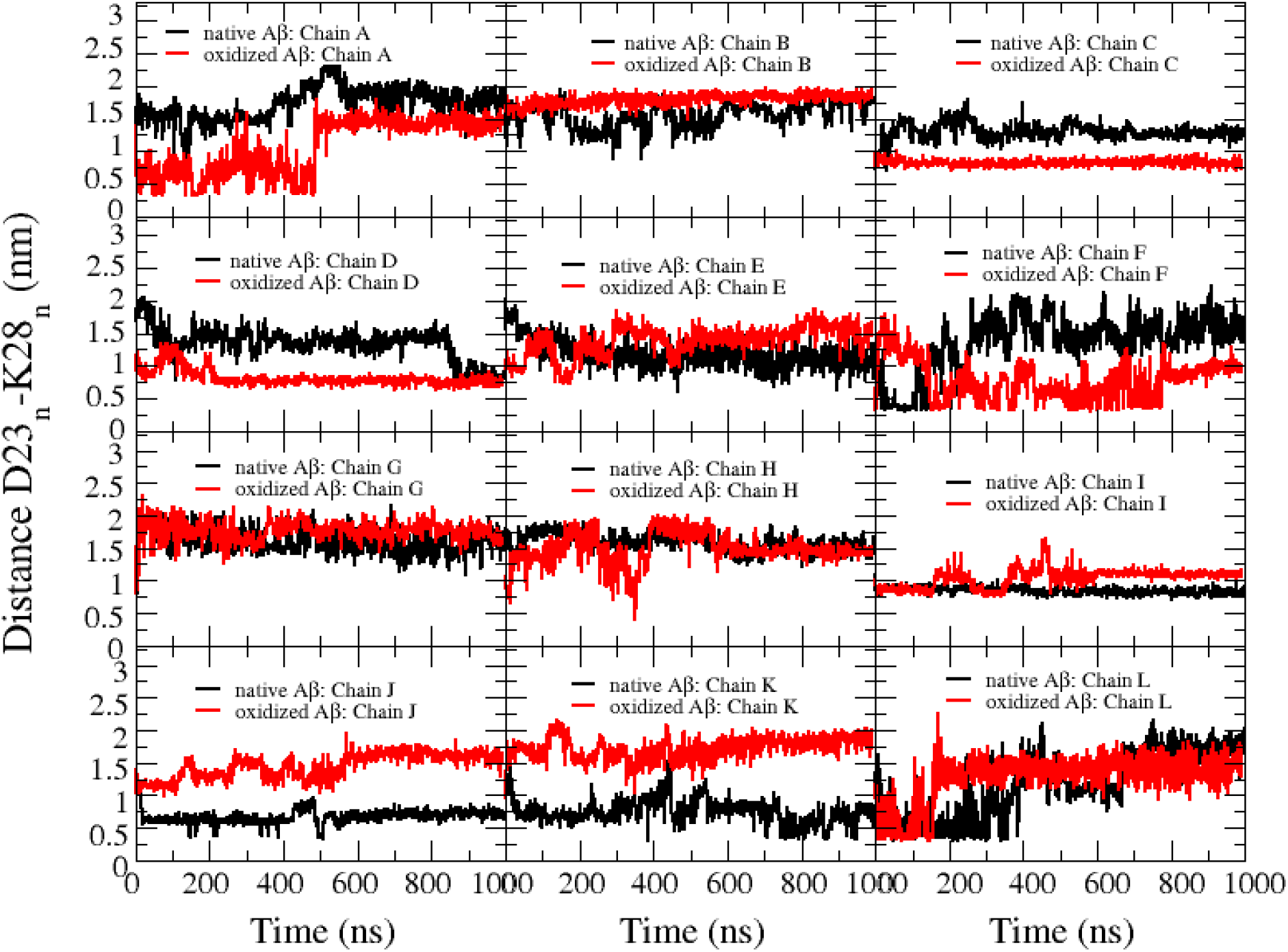
Profiles of the intra-chain Asp23-Lys28 (D23-K28) salt bridge distances of each Aβ chain (A to L) in the native (black) and oxidized (red) protofibrils for the 1 μs simulations. Subscript n in the y-axis label denotes that both D23 and K28 are from the same peptide chain.

Additionally, we also analyzed the inter-peptide salt bridges that are formed by Asp23 and Lys28 on adjacent Aβ peptides for the 1 μs simulation; results are shown in Figure 4S in Supplemental Information. These salt bridges also stabilize the peptides’ U-shape conformation and contributes to the rigidity of Aβ protofibrils and fibrils. As shown in Figure 4S, no consistent trend emerged in the comparisons of inter-peptide salt bridge distances between the oxidized and native protofibrils. Some inter-chain salt bridge distances were decreased by Met35 oxidation (chains C and D, G and H, and D and K), some remained unchanged (chains B and C, D and K, I and J), and in one neighboring pair, increased (chains J and K).

Taken together, our analysis of intra- and inter-salt bridge distances showed that oxidation did not have a significantly destabilizing effect on the turn region of the Aβ peptides’ U-shape motif, nor did it destabilize the stacking of peptides in the protofibril axial direction in each of the hexamers. Previous MD simulation studies have reported the destabilization of inter- and intrapeptide D23-K28 salt bridges by the binding of ligands such as caffeine^82^, brazilin^83^, and a resveratrol derivative.^84^ Experimental studies of the binding of these ligands to Aβ fibrils and protofibrils showed near-complete disintegration and remodeling of the Aβ fibril morphology^83,84^. In contrast, photo-oxidation of Aβ fibrils that results in Met35 oxidation has been observed to break down fibrils into smaller fibrils that retained their cross-β-sheet structure.^43^ Our simulation results are thus consistent with the experimental finding that the intra- and inter-peptide β-sheet structures that form the backbone of the fibrils were not completed disrupted by Met35 photooxidation.

### The number of hydrogen bonds in protofibrils

Experimental and theoretical investigations have shown that the extended β-sheet conformation of the Aβ protofibril and fibril are stabilized by a network of H-bonds, both intra- and inter-peptide. ^80,85^ The U-shape motif of each Aβ peptide in the protofibril is stabilized by H-bonds that form between the two β-strands. The U-shape peptides then stack axially *via* hydrogen bonds to form the extended inter-peptide β-sheet structure of the fibrils and protofibrils, with the hydrophilic surface composed of N-terminal amino acids facing the solvent and the hydrophobic surface composed of C-terminal amino acids facing the interior of the fibril and protofibril core.^86–88^ To assess the effect of Met35 oxidation on the H-bond network of the protofibril, we calculated the number of H-bonds in the native and oxidized protofibrils as a reduction in the number of H-bonds can potentially disintegrate the β-sheets, leading to the destabilization of the Aβ protofibril structure.

As shown in Figure 9, both the short and long simulations indicate that the oxidized protofibril has less H-bonds compared to the native protofibril. The difference is more pronounced in the longer 1 μs simulations where the native protofibril has an average of 161 H-bonds compared to an average of 135 H-bonds in the oxidized protofibril. Oxidation of the Met35 residues in the core of the protofibril thus caused about 16% reduction in the extended H-bond network in the protofibril.

**Figure 9.**
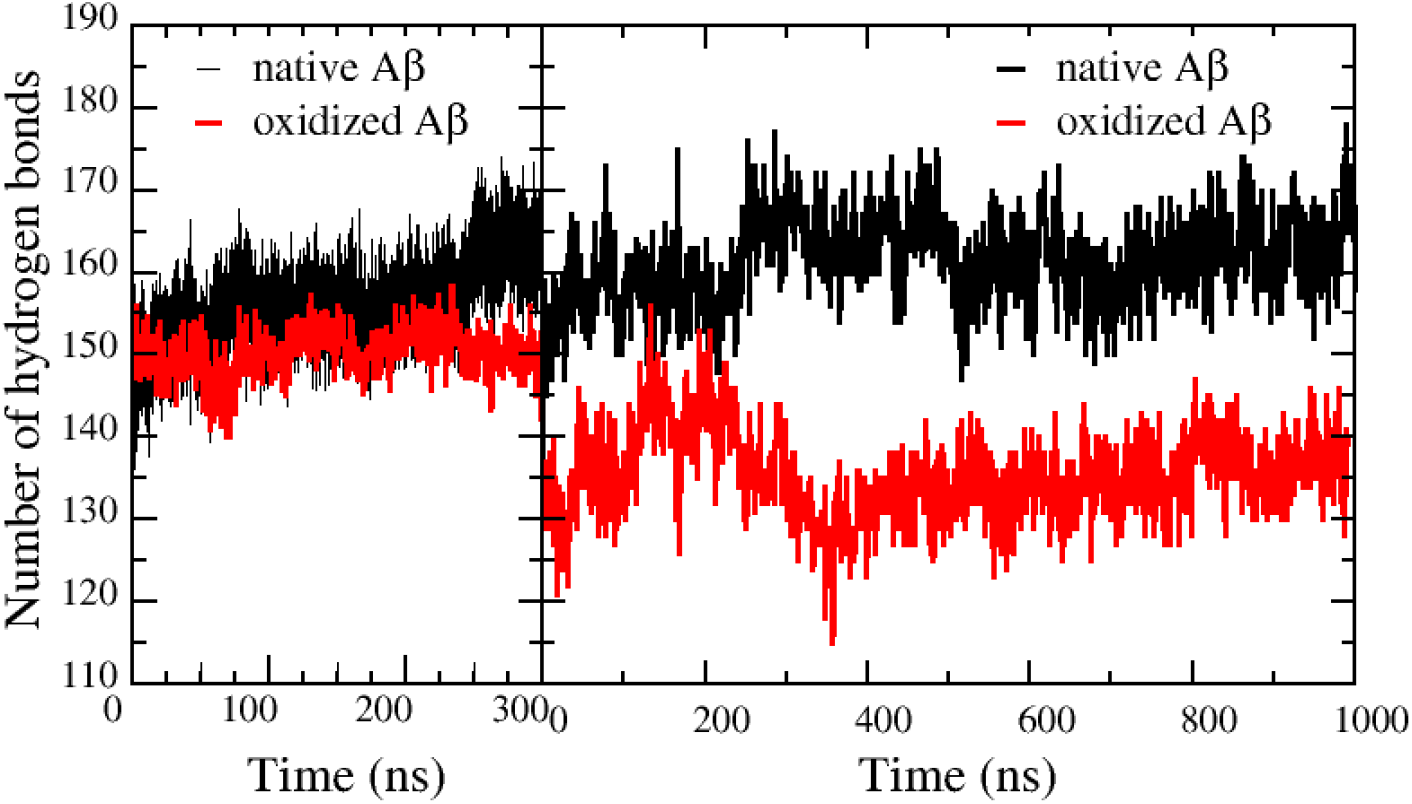
The number of hydrogen bonds in the native protofibril (black) and the Met35 oxidized protofibrils (red) for the 300 ns (left) and 1 μs (right) simulations. For the 300 ns simulations, the average number of hydrogen bonds from 3 simulation trajectories are plotted.

### Secondary structures of Aβ protofibrils

The Aβ protofibril adopts a highly ordered conformation wherein the peptides primarily form intra- and inter-molecular β-sheets. Secondary structural analysis thus gives us insights into the effect of Met35 oxidation on the structural stability of the protofibril. We employed the DSSP method^73^ to map the secondary structures of both native and oxidized protofibrils for the 1 μs simulations (Figure 10A and 10B). Secondary structure plots were also constructed from the analysis (Figure 10C and 10D).

**Figure 10.**
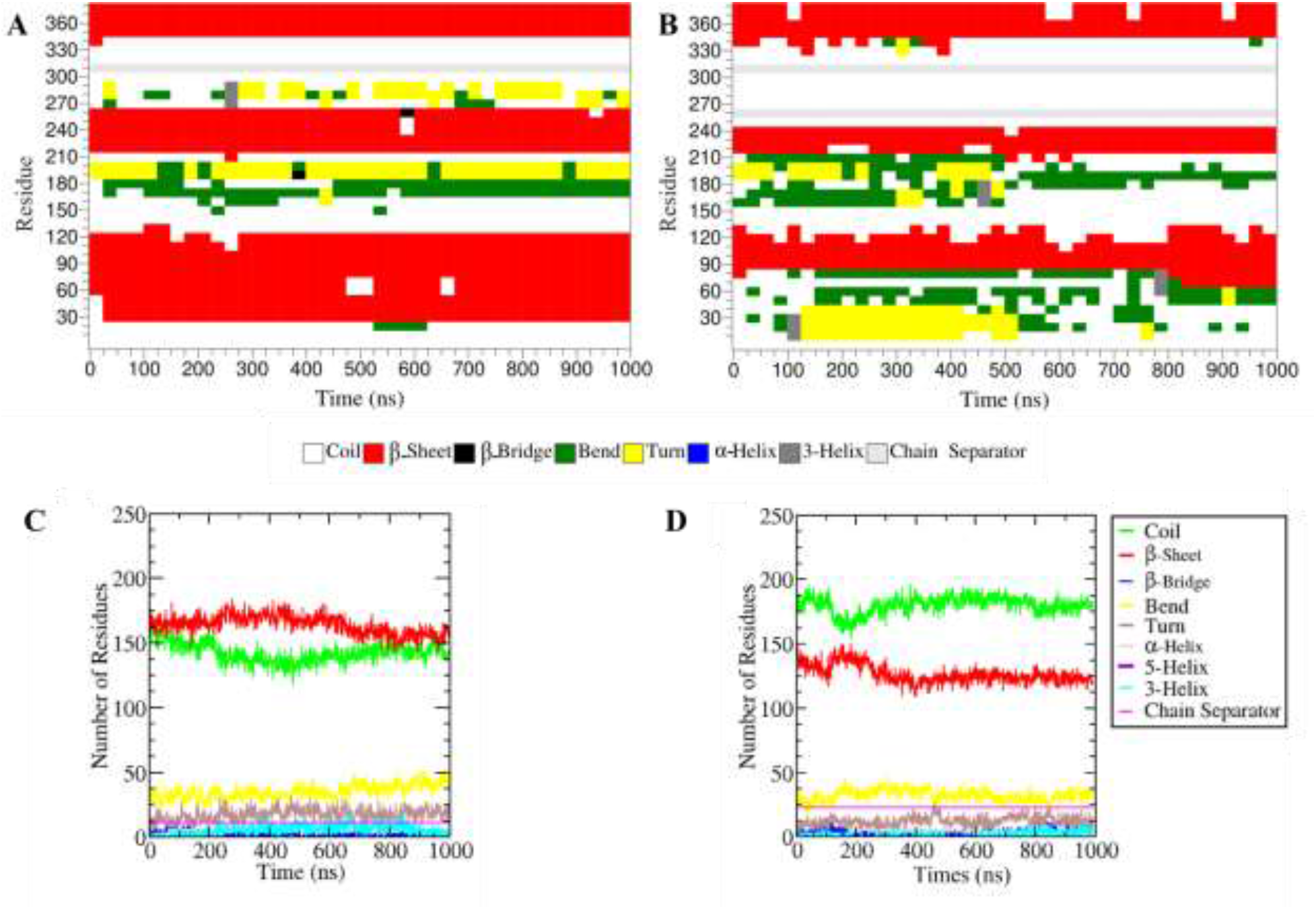
Secondary structure analysis of native and oxidized Aβ protofibrils for the 1 μs simulations. DSSP mapping of native (**A**) and oxidized (**B**) Aβ protofibrils show the secondary structures adapted by each amino acid of the 12 Aβ peptides that form the protofibril. Secondary structure plots of native (**C**) and oxidized (**D**) Aβ protofibrils show the number of amino acids that adopt each of the secondary structural elements.

As shown in Figure 10A, the predominate secondary structures in the native protofibril were β-sheets (red) and coils (white), following by turns (yellow) and bends (green). In the oxidized protofibril, the level of β-sheets was reduced, accompanied by a notable increase in the level of coils (Figure 10B). These changes are more easily visualized in plots of secondary structures of the native (Figure 10C) and oxidized (Figure 10D) protofibrils. Note that the β-sheet content decreased from an average of 164 residues in the native protofibril over the 1 μs simulation to 125 residues in the oxidized protofibril. The loss of about 40 amino acids in β-sheets is largely accounted for by a gain of about 30 residues in turns. Combined with the observation that the other secondary structures remained largely at the same low levels in the oxidized fibrils, the phototoxidation of Met35 appeared to mainly have a disruptive effect on some β-sheets (about 24%) in the protofibril and did not cause the significant formation of other, ordered secondary structures such as α-helices. Combined with results from SASA and salt bridge distances analyses, the loss of β-sheets likely occurred mainly at the hydrophobic interface between the two hexamers, and not the β-strand stacking within the hexamers.

## Discussion

Photosensitizer-induced oxidation of Aβ aggregates is being explored as a promising therapeutic strategy for the targeted degradation and clearance of the aggregates. Photo-oxidized fibrils exhibit lower toxicity *in vitro* and importantly, photo-oxidation has been reported to reduce brain Aβ aggregate levels and extend the longevity of AD animal models.^28,38,39^ The mechanism by which photo-oxidation mediates Aβ aggregate toxicity and clearance remains to be elucidated. We have shown in a recent *in vitro* study that a fibril selective photosensitizer caused clumps of Aβ40 fibrils to dissociate and fragment into smaller fibrils with light irradiation.^43^ Moreover, the oxidized fibrils retained a significant amount of β-sheet structures of the native fibrils and the ability to seed the aggregation of Aβ monomers. This partial fibril destabilization may be advantageous since more complete degradation of amyloid fibrils can potentially result in oligomers that are more toxic compared to monomeric or fibrillar Aβ conformers.^89,90^ Reduced brain aggregate levels of Aβ demonstrated in animal models may occur by the targeting and clearance of photo-oxidized aggregates through endogenous protein degradation pathways. In order to better understand photo-sensitized fibril degradation and clearance, and further develop PDT to treat AD, a detailed understanding of the effect of photo-oxidation on fibril structure and stability is needed.

Chemically, photo-sensitizers induce the oxidation of His13, His14, and Met35 residues in Aβ40 fibrils.^18,43,46^ In this study, we performed all-atom MD simulations to investigate the effect of Met35 oxidation on fibril structural dynamics and stability. Simulations up to 1 μs were carried out on an Aβ_9-40_ protofibril. This protofibril is composed of two stacked hexamers and was chosen because it contains the extended β-sheet structure that is characteristic of polymorphic Aβ fibrils. Simulations of a native protofibril containing Met35 and an oxidized protofibril where Met35 residues were replaced with methionine sulfoxide were carried out and analyzed.

Simulation snapshots show that the oxidized protofibril retained its aggregated structure (Figure 1). Twisting of Aβ chains along protofibril axis and some loss of β-sheet contacts were observed in both native and oxidized protofibrils. However, chain twisting was observed more frequently and at a higher degree in the oxidized protofibril compared to the native protofibril. β-sheet loss is also more apparent in the oxidized protofibril. Analyses of the global conformational states of the native and oxidized protofibrils indicate that Met35 oxidation has a destabilizing effect on the highly ordered and compact protofibril structure. Compared to the native protofibril, oxidized protofibril showed an approximate 30% increase in backbone Cα-RMSD, 5-14% increase in SASA, and a smaller around 3% increase in R_*g*_. In addition, 8 out of 12 chains of the oxidized protofibril showed higher residue RMSF values compared to the native protofibril, indicating that many residues in the oxidized protofibril exhibit higher flexibility.

Further analysis of the specific interactions that stabilize the extended-β-sheet protofibril conformation shows that although the oxidized protofibril contains about 16% less H-bonds and 24% less β-sheets, the intra-chain salt bridges that stabilize the U-shape of each peptide and the inter-chain salt bridges stabilize the stacking of the peptides in each of the hexamers are largely unperturbed. The intra-chain Asp23-Lys28 salt bridge that stabilizes the loop region connecting the two β-strands of the peptide U-shape is a highly conserved conformation of Aβ fibril polymorphs.^76^ The inter-chain Asp23-Lys28 salt bridge is strong and also helps to stabilize the U-shaped peptide conformation and account for the high structural rigidity of Aβ fibrils.^76^ Met35 photo-oxidation thus did not exhibit a consistent destabilizing trend in the U-shape of the peptides or the stacking of the peptides in the hexamers. It did, however, disrupt the hydrophobic interactions between the two hexamers as the Met35^ox^-Met35^ox^ distances are larger for the interior, most hydrophobic, chains compared to Met35-Met35 distances. This finding is also consistent with increases in SASA and R_*g*_ values of the oxidized protofibril.

Our MD simulation results thus indicate that the oxidation of Met35 caused some destabilization to the overall conformation of the protofibril. Specifically, Met35 oxidation that resulted in the addition of a hydrophilic oxygen disrupted the hydrophobic interface that stabilizes the stacking of the two hexamers. The oxidized protofibril is more solvent exposed, less compact, and exhibit more backbone flexibility, but retained the underlying U-shaped structure of each peptide. Although more twisting of the peptides along the protofibril axis was observed, the stacking of the peptides in the hexamers remained with Met35 oxidation. Our simulation results are thus consistent with experimental observations that photo-oxidation of Aβ40 fibril results in the dis-agglomeration and fragmentation of Aβ fibrils, but did not cause complete disruption of the fibrillar morphology or β-sheet structures.^43^

Our photo-oxidation results that show partial destabilization of preformed Aβ fibrils are different than those where Aβ fibrils were oxidized by chemical oxidants (e.g., oxidation of Aβ_1-42_ fibrils by 7.5 wt% of H_2_O_2_ that caused remodeling of fibrillar morphology to irregularly shaped rope-like structures and globules^91^ or where Aβ fibrils are destabilized by the binding of ligands (e.g., wine-related polyphenols that completely disaggregated pre-formed Aβ fibrils into disordered monomers *in vitro*.^92^ MD simulation of the binding of a related polyphenol caffeine to an Aβ_17-42_ pentamer showed destabilization of the pentamer conformation, loss of H-bonds and β-sheet structures, as well as salt bridges.^82^ This more complete disruption of the Aβ pentamer is consistent with experimental results that showed complete disaggregation of Aβ fibrils. A recent MD simulation study examined the effects of oxidation of five different residues (Met35, Phe19, Ph20, Lys16, Lys28) on the stability of an Aβ_11-42_ pentamer *via* umbrella sampling.^93^ This particular oxidation pattern was experimentally achieved by a pulsed radio-frequency cold atmospheric plasma jet that caused the complete disintegration of Aβ_1-42_ fibrils.^94^ The high level of oxidation was found in the simulation study to disrupt salt bridges and cause significant disturbance to the pentamer structure.^93^ This simulation study also showed that a low and moderate degree of oxidation (one (Met35) or three (Met35, Phe19, and Phe20) oxidized amino acids) had insignificant impact on the pentamer conformation and did not disrupt salt bridges, which is consistent with our findings in this MD study. However, we note that photosensitized oxidation also leads to the oxygenation of two histidine residues (His13 and His14) and their effects are not included in this computational study. The addition of hydrophilic oxygens to the imidazole ring of histidine can also disrupt their hydrophobic interactions and contribute to fibril destabilization.

## Conclusions

We studied the effects of Met35 oxidation on the conformation and stability of an Aβ_9-40_ protofibril, employing all atom MD simulations for up to 1 μs. The results demonstrate that the oxidation of Met35 caused some destabilization to the overall conformation of the β-sheet rich protofibril, as evidenced in increases in RMSD, R_*g*_, SASA, and RMSF values. The oxidized protofibril is thus more solvent exposed, less compact, and exhibit more backbone flexibility, which may be contributed by the destabilization of the hydrophobic interface that stabilize the stacking of two hexamers in the protofibril as evidenced by increased methionine-methionine distances in the oxidized protofibril. However, Met35 oxidation did not significantly perturb the intra- and inter-chain salt bridges that stabilize the secondary U-shaped conformation adopted by each chain nor the stacking of the peptides that form each of the hexamers. These simulation results are consistent with experimental findings that photo-oxidation caused partial destabilization of Aβ40 fibrils and did not completely disrupt the conformation and underlying secondary structures of the Aβ aggregate. This is in contrast with oxidation caused by chemical oxidants or strong oxidizing sources such as cold atmospheric plasma. This computational study thus provides molecular level insights into the partial, and more nuanced, perturbations of Aβ40 aggregates by photosensitizer induced oxidation. Combined with *in vivo* studies that demonstrated the efficacy of PDT in lowering aggregate levels and reducing neurotoxicity of Aβ aggregates in AD animal models, this investigation contributes to our future development of photo-active platforms for treating protein misfolding diseases such as AD.

## Author Contributions

F.M., T.D.M., and E.Y.C designed research; F.M. performed research and analyzed results; F.M and E.Y.C wrote the paper.

## Acknowledgements

The Extreme Science and Engineering Discovery Environment (XSEDE), which is supported by National Science Foundation grant ACI-1053575, was used for performing simulations.

## Abbreviations

Aβ: amyloid β
AD: Alzheimer’s disease
MD: molecular dynamics
PDT: photodynamic therapy
ROS: reactive oxygen species
NMR: nuclear magnetic resonance
His: Histidine
Tyr: Tyrosine
Met: Methionine
PDB: Protein Data Bank
NVT: isothermal ensemble
NPT: isobaric ensemble
LINCS: A Linear Constraint Solver for Molecular Simulations
PME: Particle Mesh Ewald
RMSD: root mean square deviation
RMSF: root mean square fluctuation
R_*g*_: radius of gyration
SASA: solvent accessible surface area
DSSP: dictionary of secondary structure of protein
XSEDE: Extreme Science and Engineering Discovery Environment

