## Supporting information for "Partial Destabilization of Amyloid-β Protofibril by Methionine Photo-oxidation: A Molecular Dynamic Simulation Study"

#### Supplemental Information

### Effect of Methionine Photo-oxidation on the Conformational Dynamics of Amyloid-β Protofibril: A Molecular Dynamic Simulation Study

Fahimeh Maghsoodi[1,2], Tye D. Martin[2], and Eva Y. Chi[2,3*]

[1]Center for Biomedical Engineering, University of New Mexico, Albuquerque, NM 87131

[2]Nanoscience and Microsystems Engineering Graduate Program, University of New Mexico, Albuquerque, NM 87131

[3]Department of Chemical and Biological Engineering, University of New Mexico, Albuquerque, NM 87131

Keywords: Alzheimer's disease, amyloid aggregates, photo-oxidation, methionine oxidation, protofibrils, stability, molecular dynamics simulation

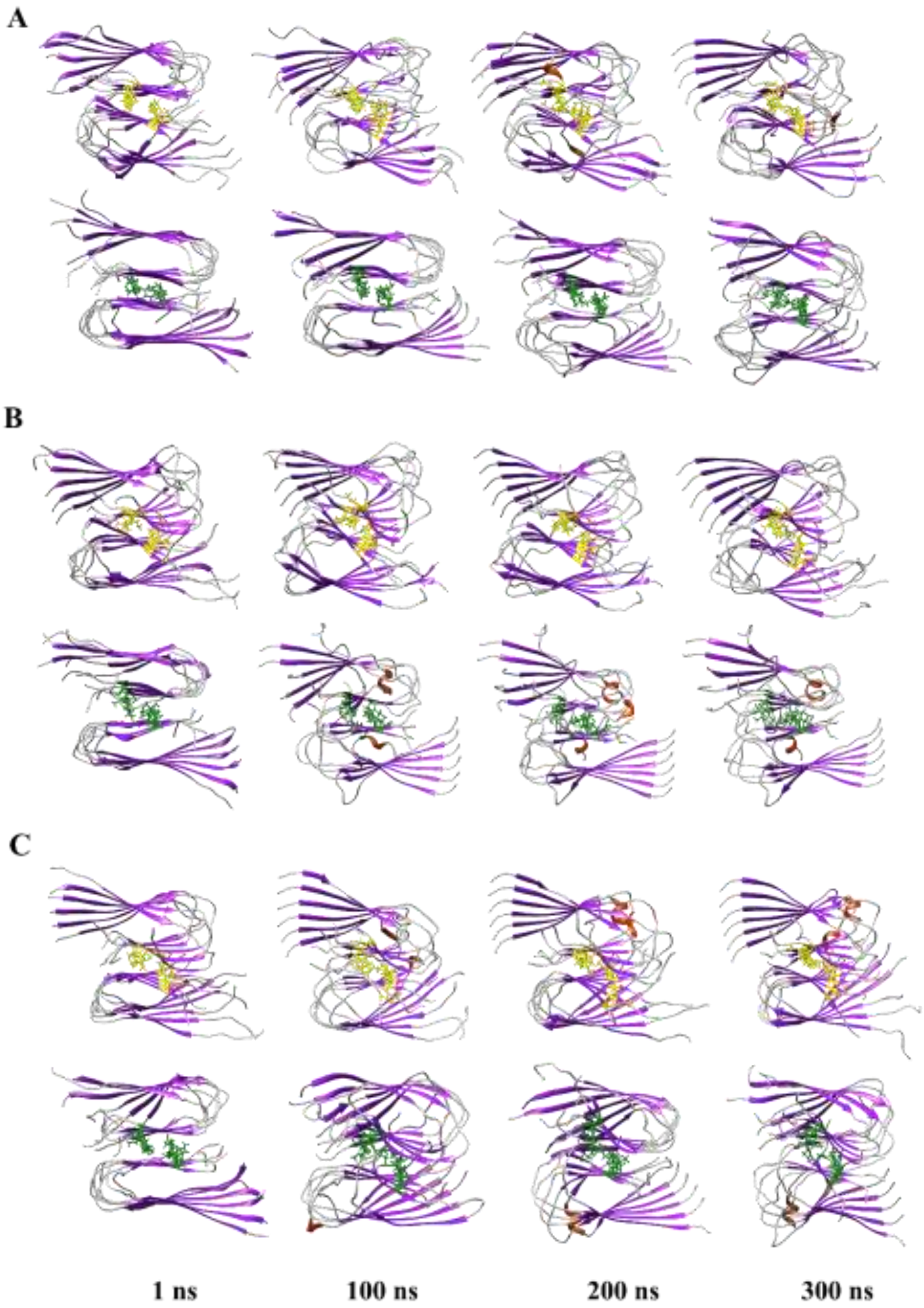


**Figure S1**. Snapshots of native and oxidized Aβ protofibrils from three 300 ns simulation trajectories (**A**, **B**, and **C**). Met35 and Met35$^{ox}$ are shown in yellow and green, respectively. β-strands, α-helices, and coils are shown in purple, orange, and grey, respectively.

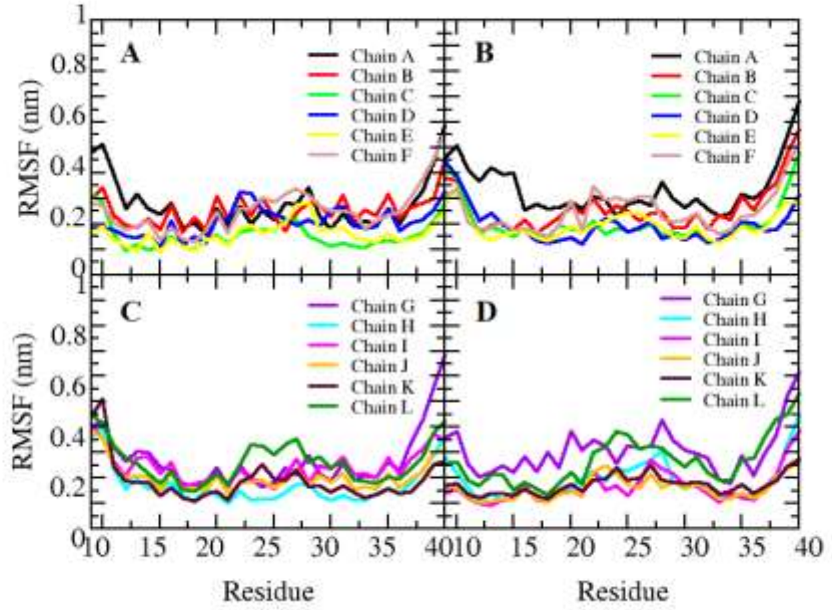


**Figure S2**. RMSF plots of different Aβ chains belonging to the top hexamer (chains A to F: **A** and **B**) and bottom hexamer (chains G to L: **C** and **D**) of the native (**A** and **C**) and oxidized (**B** and **D**) protofibrils after 300 ns of simulation. Values are averages from 3 simulation trajectories.

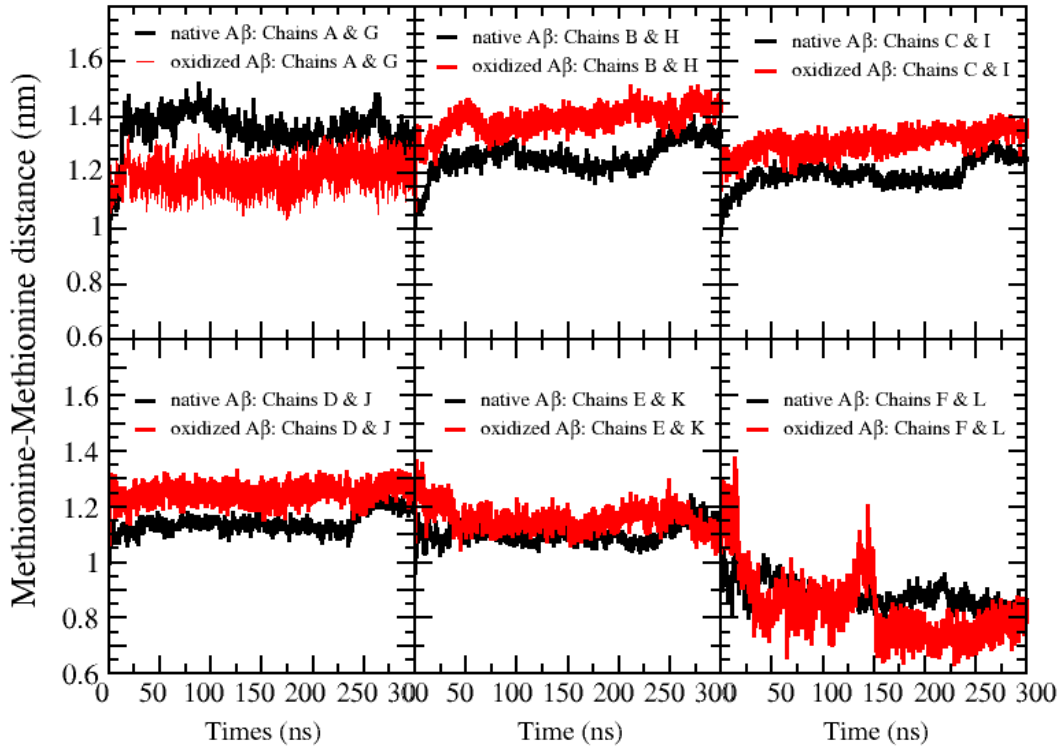


**Figure S3**. Distances between Met35-Met35 (black) and Met35ox-Met35ox (red) residues of 6 opposing Aβ chain pairs (A and G, B and H, C and I, D and J, E and K, and F and L) in the protofibrils for the 300 ns simulations. Values are averages from 3 trajectories.

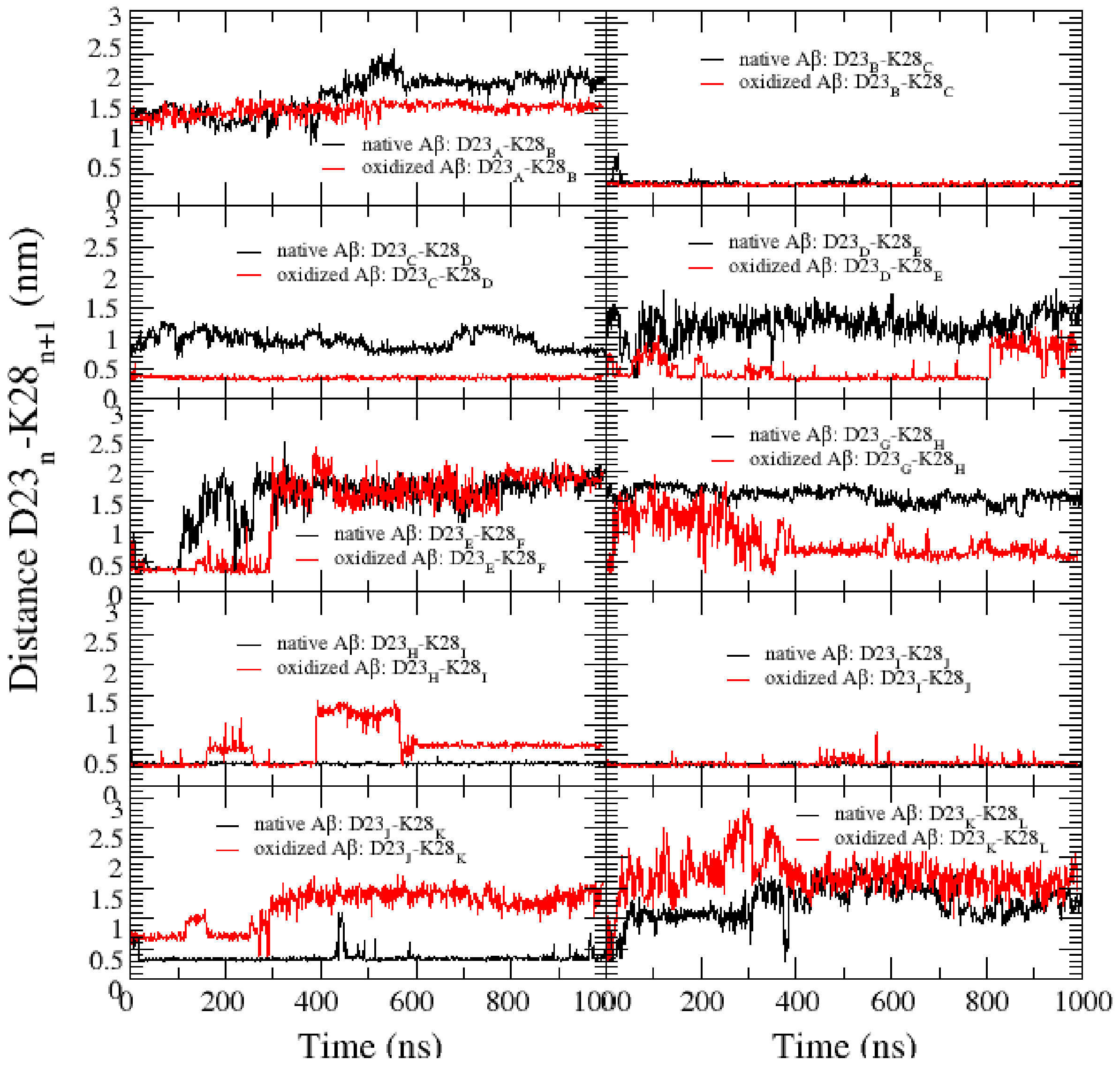


**Figure S4**. Distances of inter-peptide salt bridges formed by Asp23 and Lys28 (D23-K28) on adjacent Aβ peptides for native (black) and oxidized protofibrils (red) for the 1 μs simulations. Subscripts n and n+1 in the y-axis label denote that the D23 and K28 are from different peptide chains.
